# Sweet and magnetic: Succession and CAZyme expression of marine bacterial communities encountering a mix of alginate and pectin particles

**DOI:** 10.1101/2020.12.08.416354

**Authors:** Carina Bunse, Hanna Koch, Sven Breider, Meinhard Simon, Matthias Wietz

## Abstract

Polysaccharide particles are an important nutrient source and microhabitat for marine bacteria. However, substrate-specific bacterial dynamics in a mixture of particle types with different polysaccharide composition, as likely occurring in natural habitats, are undescribed. Here, we studied the composition, functional diversity and gene expression of marine bacterial communities encountering a mix of alginate and pectin particles. Communities were collected above macroalgal forests near Helgoland Island − where polysaccharide-rich particles might regularly occur − and exposed to a mix of magnetic particles of each polysaccharide, allowing the targeted evaluation by particle type. Amplicon, metagenome and metatranscriptome sequencing revealed that particle-associated (PA) and free-living (FL) communities significantly differed in composition and metabolism, whereas dynamics on alginate and pectin particles were unexpectedly similar. Amplicon sequence variants (ASVs) from *Tenacibaculum*, *Colwellia*, *Psychrobium* and *Psychromonas* dominated the community on both particle types. Corresponding metagenome-assembled genomes (MAGs) expressed diverse alginate lyases, several co-localized in polysaccharide utilization loci. One low-abundance MAG related to *Catenovulum* showed pectin specialization through upregulated GH53 and GH105 genes. A single *Glaciecola* ASV dominated the FL fraction, likely persisting on particle-derived oligomers through different glycoside hydrolases. The bacterial preference for alginate, whereas pectin mainly served as colonization scaffold, illuminates substrate-driven microbial dynamics within mixed polysaccharide resources. Moreover, elevated ammonium metabolism signifies nitrogen availability as important factor on particles, whereas elevated methylcitrate and glyoxylate cycles illustrate nutrient-limited conditions in the surrounding water. These insights expand our understanding of bacterial microscale ecology, niche specialization and the biological carbon pump in macroalgae-rich habitats.

## INTRODUCTION

Polysaccharides produced by marine macroalgae and phytoplankton are important ecological and biogeochemical agents, being structural and storage components for the algae as well as nutrient source for heterotrophic bacteria [1–3]. A considerable fraction of algal polysaccharides is bound in particles, hotspots of microbial activity with central implications for the biological carbon pump [4–6]. Hydrogels and transparent exopolymer particles (TEP), a subset of polysaccharide particles resulting from self-assembly of anionic polysaccharides in seawater, constitute a global amount of ~70 gigatons and are indispensable for the study of particle-microbe interactions [6–8]. The building blocks of marine hydrogels largely originate from macroalgae, in which anionic gelling polysaccharides such as alginate can constitute >50% of the biomass [9]. Natural processes of decay or exudation, such as the release of alginate and rhamnogalacturonan from widespread macroalgae [10], presumably result in formation of hydrogel scaffolds that represent hotspots for microbial life. These events potentially play an ecological role at rocky coasts of temperate seas, which harbor dense forests of macroalgae.

The chemical and structural complexity of marine hydrogels challenges the identification of specific particle-microbe relationships. To reduce this complexity, exposing synthetic model particles to natural bacterioplankton helps understanding the dynamics and drivers of particle colonization. Such approaches have identified hydrogels and other polysaccharide particles as active microbial microhabitats, which harbor distinct communities compared to the surrounding water [11–14]. Furthermore, attached microbes can undergo a temporal succession of primary degraders and opportunistic taxa [13, 15, 16]. Associated patterns in the diversity of carbohydrate-active enzymes (CAZymes), foremost polysaccharide lyases (PLs) and glycoside hydrolases (GHs), discriminate bacteria into primary degraders, secondary consumers and non-hydrolyzers [17]. The genomic co-localization of CAZymes in polysaccharide utilization loci (PUL) is thereby a common indicator of hydrolytic capacities [18].

Nonetheless, it remains enigmatic how bacterial particle utilization proceeds within the natural “particlescape” − presumably containing a mixture of particle types with different polysaccharide composition − and how these processes are shaped by the taxonomic and functional diversity of the ambient microbiota. The co-availability of hydrogels with different polysaccharide composition might initiate a segregation of bacterial populations by substrate preferences, comparable to hydrolyzing model isolates [10, 19]. In this context, the CAZyme repertoire is considered a stronger driver of niche specialization than phylogenetic relationships [17, 20]. The colonization and utilization of particle resources might also include interactions with free-living microbes, which might benefit from cross-feeding on oligosaccharides released into the water column. In addition, competition, cooperation and differing growth rates likely result in successional patterns and specific interactions [6, 21, 22].

To evaluate particle-specific bacterial dynamics in a mixture of hydrogels, we co-exposed alginate and pectin particles to bacterioplankton communities from Helgoland, a North Sea island surrounded by dense macroalgal forests [23, 24]. Due to the gelling capacities of alginate and pectin and their demonstrated release from Helgoland macroalgae [10], we assume that related particles occur in this habitat and constitute microhabitats for specialized microbiota. The co-incubation of magnetic and non-magnetic particles of each polysaccharide allowed deciphering community composition, functional potential and gene expression depending on particle type and in relation to the free-living fraction. Opposed to our hypothesis that alginate and pectin particles are utilized by different members of the ambient community, we observed similar compositional and functional patterns with predominant expression of alginate lyases. The identification of alginate as preferred substrate, whereas pectin primarily served as colonization scaffold, illuminates bacterial microhabitat ecology and carbon cycling in macroalgae-rich habitats with diverse polysaccharide budgets.

## METHODS

### Characteristics of polysaccharide particles

Custom polysaccharide particles consisting of alginate (CAS #9005-38-3) or pectin (CAS #9000-69-5) were fabricated by geniaLab (Braunschweig, Germany) in four different versions: magnetic alginate and pectin particles (polysaccharide coated on magnetite core) as well as non-magnetic alginate and pectin particles (polysaccharide coated on ferrous iron core). Particles contained on average 4% polysaccharide (Supplementary Methods), approximating the 1:100 solid:solvent ratio in natural hydrogels [8]. Magnetic and non-magnetic particles of each polysaccharide were prepared to allow co-incubation of both polysaccharides and hence provide similar selection pressures per treatment (i.e. always two available particle types, only varying in magnetism). Applying external magnetic force (Supplementary File 1) allowed the targeted and sterile sampling of particle types.

### Seawater sampling and experimental set-up

Seawater was sampled from approx. 1 m depth above macroalgal forests at Helgoland Island (54°11’26“N 7°52’00”E) in June 2017. Seawater was filtered through a 100 μm mesh, brought to the lab within two hours, and filtered again through a 20 μm mesh. Each 12 L of filtered seawater were distributed into 20 L Clearboy bottles (Nalgene, Rochester, NY) previously rinsed with the same seawater. Per bottle, 10 mM NaNO_3_ and 1 mM NaH_2_PO_4_ were added to counteract nitrogen or phosphorus limitation. Three experiments were set up in triplicate: (i) magnetic alginate particles and non-magnetic pectin particles, (ii) non-magnetic alginate particles and magnetic pectin particles, and (iii) control without particles (Supplementary Fig. 1). Each particle type was added at 3500 L^−1^, resulting in ~42,000 particles per bottle. Bottles were incubated statically at 15°C (approx. *in situ* temperature) in the dark.

### Sampling and nucleic acid extraction of particle-associated and free-living cells

250 mL of the original seawater were filtered onto 0.2 μm polycarbonate filters for determination of the ambient bacterial community (T0). Filters were flash-frozen in liquid nitrogen and stored at ‒80°C. Incubations were sampled after 24, 48 and 60h. At each sampling point, bottles were mixed by inversion and ca. 550 mL withdrawn into rinsed measuring cylinders. Each sample was aliquoted (2×250 mL) into sterile RNase-free Nunc tubes (cat. no. 376814; Thermo Fisher Scientific, Waltham, MA). Particle-associated communities on magnetic alginate (AlgP) or pectin particles (PecP) were sampled by holding a neodymium magnet (cat. no. Q-40-10-10-N; Supermagnete, Gottmadingen, Germany) next to the tube. AlgP and PecP were washed with sterile seawater (filtered through 100 μm and 20 μm; mixed 3:1 with ddH_2_O to prevent salt precipitation during autoclaving for 20 min at 121°C) and transferred to 2 mL RNase-free microcentrifuge tubes. The supernatant was transferred to a separate tube and non-magnetic particles were removed by filtration through 5 μm polycarbonate filters. The flow-through was captured on 0.2 μm polycarbonate filters to obtain the free-living community (FL). All samples were directly flash-frozen in liquid nitrogen and stored at ‒80 °C. Simultaneous extraction of DNA and RNA was done using a modification of [39]. Purified DNA and RNA were sent on dry ice to DNASense (Aalborg, Denmark) for quality control and sequencing. For particles, several subsamples per replicate were pooled to obtain sufficient DNA and RNA (Supplementary Table 1).

### 16S rRNA gene amplicon sequencing

Briefly, the V3–V4 region of bacterial 16S rRNA genes was sequenced using primers 341F-806R [25] with MiSeq technology (Illumina, San Diego, CA). Internal company standards worked as expected (Supplementary Methods). Reads were classified into amplicon sequence variants (ASVs) using DADA2 [26] and taxonomically assigned using SILVA v132 [27]. Rarefaction analysis showed that diversity was reasonably covered (Supplementary Fig. 2). Replicates were congruent per treatment and time, without significant differences in Bray-Curtis dissimilarities (PERMANOVA; *p* = 0.72 to 0.98). Furthermore, FL communities from AlgP and PecP were congruent as expected, and FL data combined in subsequent analyses.

### Metagenomics

As amplicon data confirmed the consistency of replicates, DNA from the three AlgP, PecP and FL replicates at 24 h and 60 h were pooled, respectively. DNA was quantified using Qubit (Thermo Fisher Scientific) and fragmented to ~550 bp using M220 using microTUBE AFA fiber screw tubes (Covaris, Woburn, MA) for 45s at 20°C with duty factor 20%, peak/displayed power 50W, and cycles/burst 200. Libraries were prepared using the NEB Next Ultra II kit (New England Biotech, Ipswich, MA) and paired-end sequenced (2×150bp) on a NextSeq system (Illumina). Adaptors were removed using cutadapt v1.10 [28] and reads assembled using SPAdes v3.7.1 [29]. Genes were predicted using Prokka [30] and assigned to KEGG categories using KAAS-SBH-GhostX [31]. CAZymes were predicted using dbCAN2 with CAZyDB v8 [32], only considering hits with >80% coverage. Ammonium transporters were predicted by BLASTp of AmtB (P69681) in the Transporter Classification Database [33].

### Metatranscriptomics

RNA was quantified in duplicate per sample using the Qubit BR RNA assay (Thermo Fisher Scientific). RNA quality and integrity were confirmed using TapeStation with RNA ScreenTape (Agilent, Santa Clara, CA). rRNA was depleted using the Ribo-Zero Magnetic kit (Illumina) and residual DNA removed using the DNase MAX kit (QIAGEN, Hilden, Germany). Following sample cleaning and concentrating using the RNeasy MinElute Cleanup kit (QIAGEN), rRNA removal was confirmed using TapeStation HS RNA ScreenTapes (Agilent). Sequencing libraries were prepared using the TruSeq Stranded Total RNA kit (Illumina), quantified using the Qubit HS DNA assay (Thermo Fisher Scientific), and size-estimated using TapeStation D1000 ScreenTapes (Agilent). For RNA from particle samples, 4-5 subsamples per replicate were pooled in equimolar concentrations and sequenced on a HiSeq2500 in a 1×50 bp Rapid Run (Illumina). As the first sequencing run did not deliver sufficient data for seven metatranscriptomes, a second run was performed on the same library. PCA confirmed consistent sequencing runs (data not shown), and read counts were subsequently aggregated. Raw fastq sequence reads were trimmed using USEARCH v10.0.2132 [34] using -fastq_filter and settings -fastq_minlen 45 -fastq_truncqual 20. rRNA reads were removed using BBDuk (http://jgi.doe.gov/data-and-tools/bb-tools) using the SILVA database as reference [27]. Reads were mapped to the predicted genes using Minimap2, discarding reads with sequence identities <0.98. Relative transcript abundances were obtained by dividing raw counts by the length of each gene (RPK) and normalized by per-million scaling factors. Resulting transcripts per million (TPM) were summed per gene annotation (Supplementary Table 2). Differential transcript abundances were calculated on raw read counts using the default DESeq2 workflow in R v3.6 [35, 36] in RStudio (https://rstudio.com), only considering log2-fold changes >2 with *padj* < 0.001 (Supplementary Table 3).

### MAG binning and analysis

Binning of metagenome-assembled genomes (MAGs) was done using MetaBat2 and mmgenome2 [37, 38]. Reads were mapped back to the assembly using Minimap2 v2.5 [39] and the average coverage of each bin calculated using mmgenome2 (weighted by scaffold sizes). Based on results from CheckM and GTDB-Tk [40, 41], we selected five near-complete MAGs (≥90% estimated genome completeness and <5% genome contamination) representing the major taxa in amplicon data (Table 1, Supplementary Table 4). Whole-genome comparison with type strains was carried out using the MiGA web application [42]. Normalized coverage in metagenomic data was calculated following [43] after multiplying the coverage of each MAG in every sample with a normalization factor (sequencing depth of the largest sample divided by the sequencing depth of each individual sample). A maximum-likelihood phylogeny based on 92 core genes identified using the UBCG pipeline [44], including medium-quality MAGs (>70% estimated completeness/<10% contamination) assigned to the same genus plus related genomes from public databases, was calculated using RaxMLHPC-Hybrid v8.2.12 with the GTRGAMMA substitution model and 1,000 bootstrap replicates on the CIPRES Science Gateway v3.3 [45, 46]. Genes were assigned to KEGG categories using KAAS, and pathways reconstructed from these predictions using KEGG Pathway Mapper [31, 47]. Gene annotations were refined using UniProtKB/Swiss-Prot [48] by custom-BLAST in Geneious v7 (https://www.geneious.com). Genes for processing alginate and pectin monomers were predicted based on the fully reconstructed pathways in *Alteromonas macleodii* and *Gramella forsetii* [10, 49]. For comparative purposes, strains B3M02 and E3R01 [15] were re-annotated with dbCAN2 v8.0 and compared to CAZymes from MAG26 and MAG73 using custom-BLAST, only considering hits with >30% query coverage and >40% amino acid identity. Average nucleotide identities (ANI) between genomes were calculated using enveomics [50]. Prophages and biosynthetic gene clusters were predicted using PHASTER and antiSMASH 5.0 respectively [51, 52]. Amino acid sequences of PL7 genes from MAG73 were aligned using MAFFT algorithm E-INS-i with default parameters [53]. A maximum-likelihood phylogeny with 500 bootstrap replicates was calculated using RaxML v8.0 [45] and the WAG+G+F substitution model determined using ModelTest-NG [54], both run on the CIPRES Science Gateway [46].

**Table 1:** Characteristics of near-complete metagenome-assembled genomes.

| MAG | Taxonomy (GTDB-tk) | Complete-<br>ness | Contami-<br>nation | Size<br>(Mbp) | Genes | CAZy-<br>mes | PLs /<br>GHs |
| --- | --- | --- | --- | --- | --- | --- | --- |
| MAG24 | *Alteromonadaceae*;<br>*Colwellia* | 90 | 3.0 | 3.27 | 2886 | 91 | 16 / 18 |
| MAG26 | *Flavobacteriaceae*;<br>*Tenacibaculum* | 97 | 2.0 | 3.06 | 2781 | 65 | 17 / 9 |
| MAG32 | *Alteromonadaceae*;<br>*Glaciecola* | 96 | 0.6 | 2.77 | 2575 | 49 | 3 / 18 |
| MAG57 | *Psychrobiaceae*;<br>*Psychrobium* | 95 | 0.9 | 2.99 | 2690 | 61 | 16 / 10 |
| MAG73 | *Psychromonadaceae*;<br>*Psychromonas* | 94 | 1.0 | 3.69 | 3337 | 110 | 22 / 34 |

### Data availability

All sequencing data has been deposited at NCBI under BioProject PRJEB38771. Annotated metagenome contigs and near-complete MAGs, protein fasta files of all genes, ASV tables and the full DNA-RNA extraction protocol are available for review under https://owncloud.mpi-bremen.de/index.php/s/QEvzjh5Fx3VYf6m, to be published in final form on https://zenodo.org upon acceptance. Major R packages for analysis and visualization included tidyverse, pheatmap, ComplexHeatmap and PNWColors [55–58].

## RESULTS AND DISCUSSION

We studied taxonomic diversity, functional capacities and gene expression of particle-attached (PA) marine bacterial communities on alginate (AlgP) and pectin particles (PecP) in comparison to their free-living (FL) counterparts. Therefore, synthetic AlgP and PecP were co-exposed to natural bacterioplankton collected near Helgoland, an island in the southern North Sea surrounded by macroalgal forests and hence considerable polysaccharide budgets (Supplementary Fig. 1A). The co-incubation of magnetic AlgP and non-magnetic PecP, and vice versa, in parallel experiments allowed the targeted separation of communities on each particle type (Supplementary Fig. 1B).

### Do AlgP, PecP and FL differ in community composition and temporal variability?

Amplicon sequencing of bacterial 16S rRNA genes revealed significant differences between PA and FL communities over an incubation period of 60 h (PERMANOVA, *p* < 0.001), but substantial overlap between AlgP and PecP (Fig. 1A, Supplementary Fig. 3A). Significant compositional differences from the ambient seawater and controls without added particles (PERMANOVA, *p* < 0.001) confirmed these observations as true biological dynamics.

**Fig. 1:**
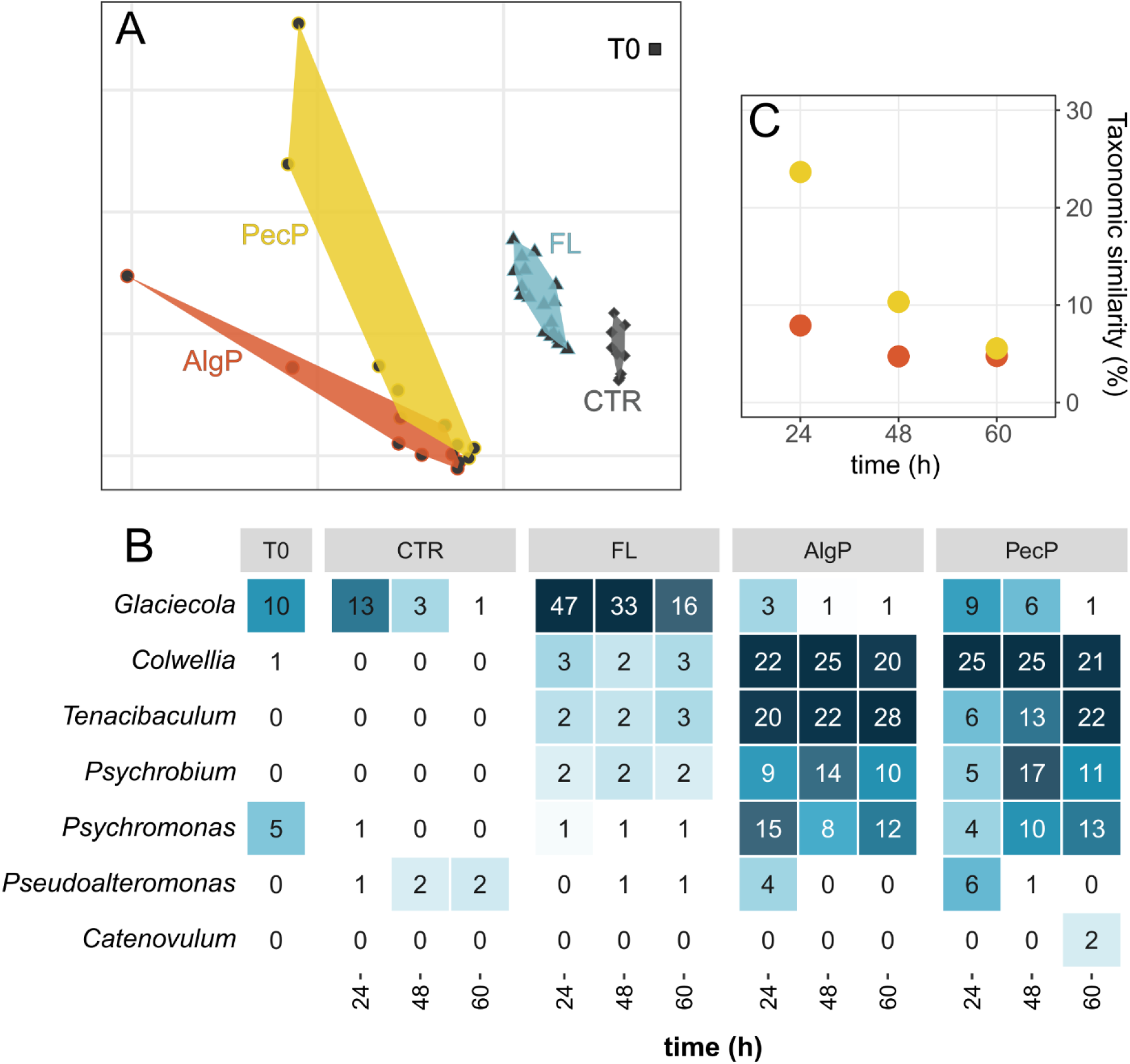
Community composition based on amplicon sequence variants. **A:** Non-metric multidimensional scaling (stress value = 0.07) illustrating marked compositional differences between particle-associated (PA), free-living (FL), control (CTR) and ambient bacterial communities (T0). **B:** Relative abundances of dominant genera on alginate (AlgP) and pectin particles (PecP) in comparison to FL, CTR and T0 communities. **C:** Taxonomic overlap based on Bray-Curtis dissimilarities in AlgP and PecP compared to T0 communities.

Amplicon sequent variants (ASVs) affiliated with *Tenacibaculum* (Bacteroidetes: Flavobacteriales), *Colwellia*, *Psychromonas* and *Psychrobium* (Gammaproteobacteria: Alteromonadales) constituted up to 60% of both AlgP and PecP communities (Fig. 1B). Hence, particle colonization largely related to few dominant taxa, comparable to other marine polysaccharide particles [13, 15]. Enrichment of these taxa compared to the FL fraction (Kruskal-Wallis test, *p* < 0.001) mirrors their occurrence on marine algae, with considerable CAZyme repertoires in some representatives [59–62]. The *Colwellia* population comprised an assemblage of diverse ASVs, appearing from nearly undetectable levels in the ambient community (Supplementary Fig. 4). This rapid stimulation on microdiverse levels indicates fast metabolic rates and high adaptability [63]. Notably, both *Colwellia* and *Tenacibaculum* can be enriched on diseased algae [64, 65] and hence under circumstances when algal polysaccharides might be released and self-assemble into particles. The ecological relevance of our observations is supported by the frequent occurrence of *Tenacibaculum* and *Psychromonas* during phytoplankton blooms near Helgoland, when bacterial dynamics are largely driven by algal carbohydrates [66–68].

*Glaciecola* (Alteromonadales) dominated the FL community (Kruskal-Wallis test; *p* < 0.0003), with an average abundance of >30% during the first 48 h with low alpha-diversity (Fig. 1B, Supplementary Fig. 3B). Notably, the population was dominated by a single ASV (Supplementary Fig. 4), suggesting that nutrient scarcity in FL favored highly competitive genotypes. This finding underlines that specific biogeochemical conditions can induce the predominance of single community members [69]. We hypothesize that *Glaciecola* largely persisted as secondary consumer of particle-derived substrates, supported by genomic evidence from the major corresponding MAG (see below).

Overall, there was little temporal variability in community composition, although we possibly missed rapid successional dynamics as observed in related studies [13, 15]. One exception was *Pseudoalteromonas*, whose sole occurrence at 24 h on both particle types (Fig. 1B; Kruskal-Wallis test, *p* = 0.002) signifies a polysaccharide pioneer [17]. This notion is supported by alginolytic and pectinolytic capacities of various *Pseudoalteromonas* species, amplified by their overall high responsiveness to nutrient input [70, 71]. The AlgP microbiota established within the first 24 h and then remained unchanged, whereas PecP were colonized more slowly and retained higher similarities to the ambient community at 24 h (Fig. 1C). Especially *Tenacibaculum* established with temporal delay, peaking after 60 h (Kruskal-Wallis test, *p* = 0.04). Overall, the substantial overlap between AlgP and PecP contradicted our hypothesis that the ambient community would segregate by particle type. Faster colonization of AlgP indicates that mainly alginate was used as substrate; these aspects are discussed below in context of metagenomic and metatranscriptomic evidence. One notable exception was *Catenovulum* (Alteromonadales), which solely established on PecP after 60 h (Fig. 1B; Kruskal-Wallis test, *p* = 0.01) and was the only taxon linked to pectin degradation (see below).

### Do AlgP, PecP and FL communities differ in functional diversity and gene expression?

As taxonomic and metagenomic richness are overall connected [72], we expected contrasting functional potentials in PA and FL communities, whereas AlgP and PecP should be largely congruent. However, AlgP and PecP might differ in gene expression patterns, as these can be independent from taxonomic composition [72]. For instance, certain taxa encode both alginate and pectate lyases [10] and might express these genes differentially depending on particle type. To evaluate these aspects, we analyzed the metagenome (24 h and 60 h) and metatranscriptome (60 h) of AlgP and PecP communities in relation to the FL fraction (Supplementary Table 1). This approach included both community-wide and genome-centric perspectives through metagenome-assembled genomes (MAGs).

The complete metagenomic library of 21 gigabases comprised ~192,000 predicted genes, of which 47% were functionally annotated using UniProtKB, KEGG and/or COG databases. Two percent of all genes were predicted to encode CAZymes according to dbCAN2 (Supplementary Table 2). Transcripts from the citrate cycle, glycolysis/gluconeogenesis and amino acid metabolism were abundant in both PA and FL metatranscriptomes, but many pathways markedly differed (Supplementary Table 3). Overall, ~60% of all transcripts were differentially abundant between PA and FL communities (Fig. 2A, Supplementary Table 3), matching metatranscriptomic evidence in other habitats [73]. Transcript abundances of glutamine synthetase, one key enzyme of bacterial nitrogen assimilation converting ammonium into glutamine, peaked in PA communities (Fig. 2B). This observation indicates considerable ammonium uptake and protein biosynthesis among PA bacteria, consistent with abundant transcripts of related transporter and regulator genes (Fig. 2B; Wilcoxon rank-sum test, *p* < 0.05). In contrast, glyoxylate, dicarboxylate and pyruvate metabolism peaked in the FL fraction (Supplementary Table 3). Induction of the methylcitrate cycle and the glyoxylate shunt (Fig. 2B; Wilcoxon rank-sum test, *p* < 0.01) is a symbol of the substrate-limited FL niche, matching transcriptomic responses of starved bacterioplankton [74]. These pathways likely promoted persistence by generating energy from short-chain fatty acids, alleviating iron limitation or oxidative stress, and overall enhancing resource exploitation [75–79]. In *Alteromonas macleodii*, similar expression patterns were interpreted as maintenance metabolism [10, 80, 81]. Notably, the biosynthesis of valine, leucine and isoleucine peaked in PA, but their degradation in FL communities (Supplementary Table 3; Wilcoxon rank-sum test, *p* < 0.01). This observation suggests cross-feeding of amino acids from actively growing PA to nutrient-limited FL bacteria. Leucine exchange has been reported between bacteria on polysaccharide particles and the surrounding water [15] and might be a stabilizing component of their interactions [82].

**Fig. 2:**
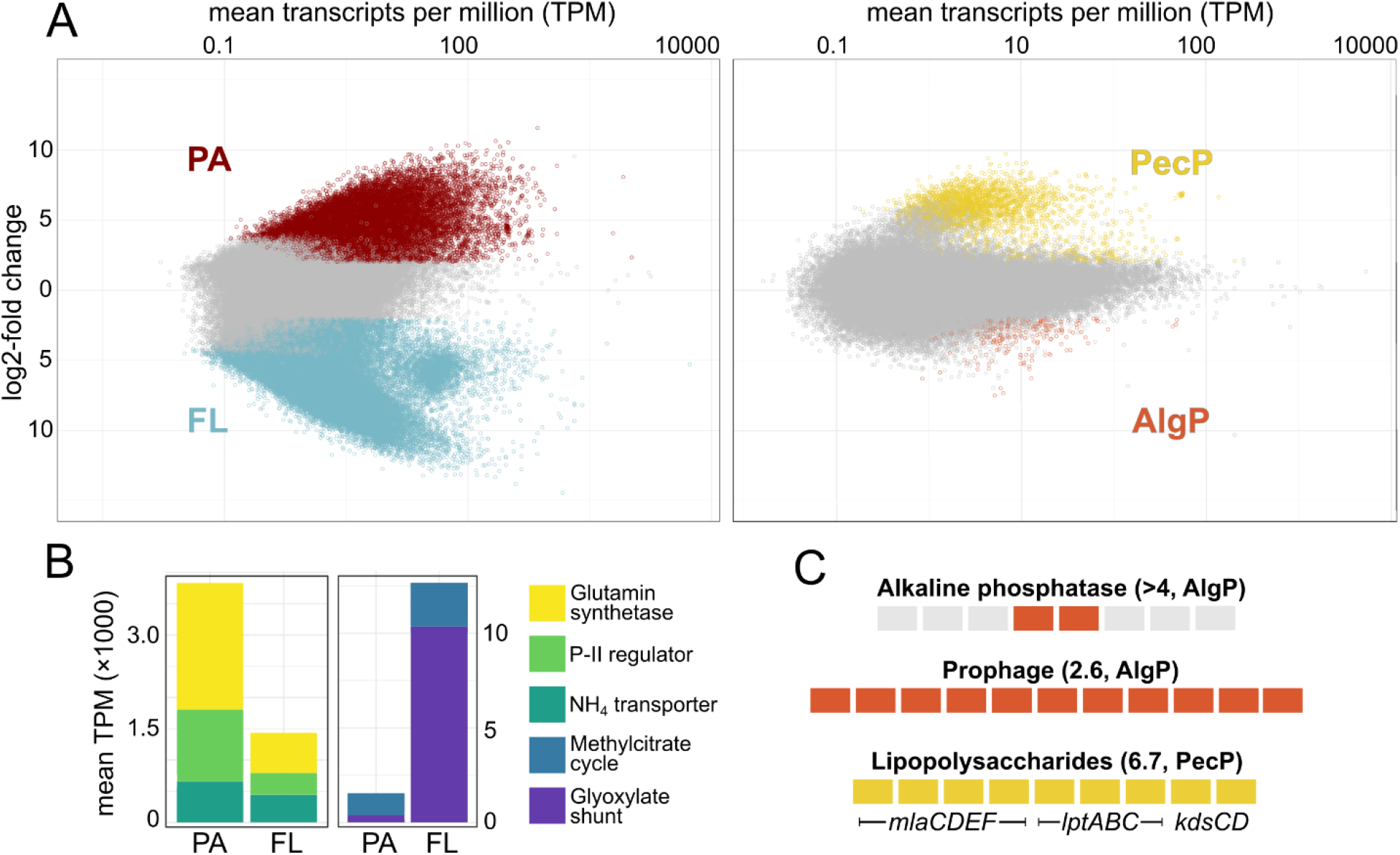
General metagenomic and metatranscriptomic patterns. **A:** Differential transcript abundances in PA versus FL (left panel) and AlgP versus PecP communities (right panel; colored dots denote DESEq2 log2-fold changes >2 with *padj* <0.001). **B:** Selected genes and pathways with higher transcript abundances in PA or FL, including glutamine synthetase (EC number 6.3.1.2) and ammonium transporters (homologs of AmtB [33]). “Methylcitrate cycle” is the sum of methyl-isocitrate lyase, methyl-aconitate isomerase, methyl-citrate synthase and aconitate hydratase genes (COG2513, COG2828, COG0372, COG1048). “Glyoxylate shunt” is the sum of isocitrate lyase and malate synthase genes (EC numbers 4.1.3.1 and 2.3.3.9). P-II regulator: Nitrogen regulatory protein P-II (COG0347). **C:** Selected gene clusters with higher transcript abundances on AlgP (orange-colored) or PecP (yellow-colored), encoding alkaline phosphatases (locus tags 41274−41275), a predicted prophage (38698−38711) and lipopolysaccharide-related *mla*, *lpt* and *kds* pathways (73039−73048). Values in parentheses designate log2-fold changes.

Communities on AlgP and PecP only slightly differed in functional potential and gene expression (Fig. 2A), compliant with their compositional overlap (Fig. 1). Only 2% of transcripts were differentially abundant, without community-wide patterns in specific functional categories (Supplementary Table 3). On AlgP, higher transcript abundances of alkaline phosphatase genes possibly counteracted beginning phosphate limitation, comparable to late stages of natural TEP colonization [83]. Furthermore, higher transcript abundances of predicted prophages (Fig. 2C, Supplementary Table 3) indicates the induction of lytic cycles and corresponding release of organic matter [84]. These events potentially stimulated secondary consumers such as *Aureispira*, which only appeared after 60 h (Kruskal-Wallis test, *p* = 0.01). This predatory taxon can feed on metabolic products or cell debris from other bacteria, fueled by its capacity to adhere to anionic polysaccharides [85]. On PecP, a single MAG affiliated with *Catenovulum* accounted for the vast majority of differentially abundant transcripts, supporting the predisposition of this taxon towards pectin (see below). For instance, the upregulation of lipopolysaccharide-related *mla*, *lpt* and *kds* genes presumably stimulated biofilm formation, an important advantage for colonization and assimilation of particulate substrates [86].

### Diversity and expression of CAZymes on community level

Similarities between AlgP and PecP extended to comparable CAZyme profiles, with a predominance of PL6 and PL7 alginate lyases on both particle types (Supplementary Table 2). PL7 genes for the initial depolymerization of alginate, approximately half including a CBM32 carbohydrate-binding domain, peaked in both copy numbers and transcript abundances (Fig. 3A, Supplementary Table 2). PL15, PL17 and PL18 genes for the processing of released oligomers were less numerous but considerably transcribed (e.g. locus tags 183566 and 114168), with the highest transcript abundance of all CAZymes in a PL18 gene (locus tags 127388). We only detected three PL1 pectate lyases and few other pectin-related genes (CE8, GH28, GH105). These results corroborate that pectin was not a prime bacterial substrate, although pectinolytic bacteria occur in diverse marine habitats [70, 87, 88] and pectinous substrates are exuded by Helgoland macroalgae [10]. Altogether, the evidence suggests that PecP primarily served as colonization scaffolds. The fast sinking of the relatively large particles (diameter ~150 μm) resulted in a loose bottom layer, with close spatial contact of both particle types. This “particlescape” potentially allowed cross-particle interactions and utilization of alginate, even if attached to PecP. Nonetheless, significantly higher abundances of alginate lyase transcripts on AlgP (Wilcoxon rank-sum test, *p* = 0.0002 to 10^−16^) indicates that PecP associates were less hydrolytic, possibly attributed to diffusion losses.

**Fig. 3:**
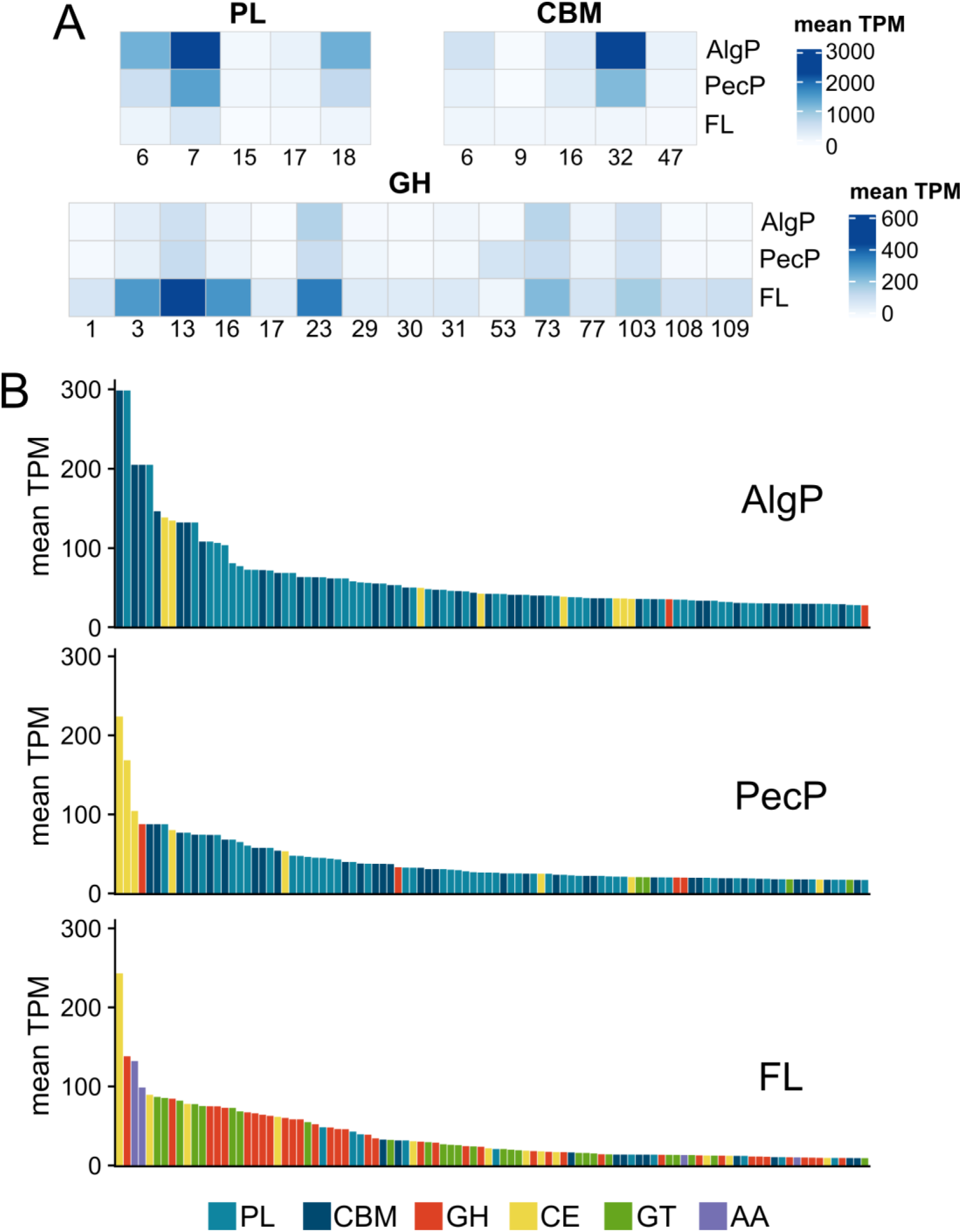
Community-wide diversity and expression of CAZymes. **A:** Average transcript abundances of polysaccharide lyases (PL), carbohydrate-binding modules (CBM) and glycoside hydrolases (GH) with mean TPM >50. **B:** Top100 CAZymes with highest expression.

The FL community showed a completely different CAZyme signature, with elevated transcript abundances of glycoside hydrolases (Fig. 3A−B). Predominance of families GH3, GH13, GH16 and GH23 (Fig. 3A) matches the hydrolase repertoire of FL bacteria during phytoplankton blooms around Helgoland [67, 89]. Hence, our observations resemble responses of natural bacterioplankton to oligosaccharide mixtures. Although these GH families are mainly associated with α-1,4-glucan, β-1,3-glucan and peptidoglycan degradation [90], our observations indicate a broader range towards oligomers of anionic polysaccharides.

### CAZymes and PUL on genomic level

Analysis of five near-complete MAGs (>90% completeness at <5% contamination) supported the genomic and ecological differentiation between PA and FL communities (Table 1, Supplementary Table 4). Core-gene phylogeny demonstrated that these MAGs represent the major PA and FL members *Colwellia*, *Tenacibaculum*, *Psychrobium, Psychromonas* and *Glaciecola* (Fig. 4A). Accordingly, their normalized coverage matched amplicon-based abundances (Fig. 4B).

**Fig. 4:**
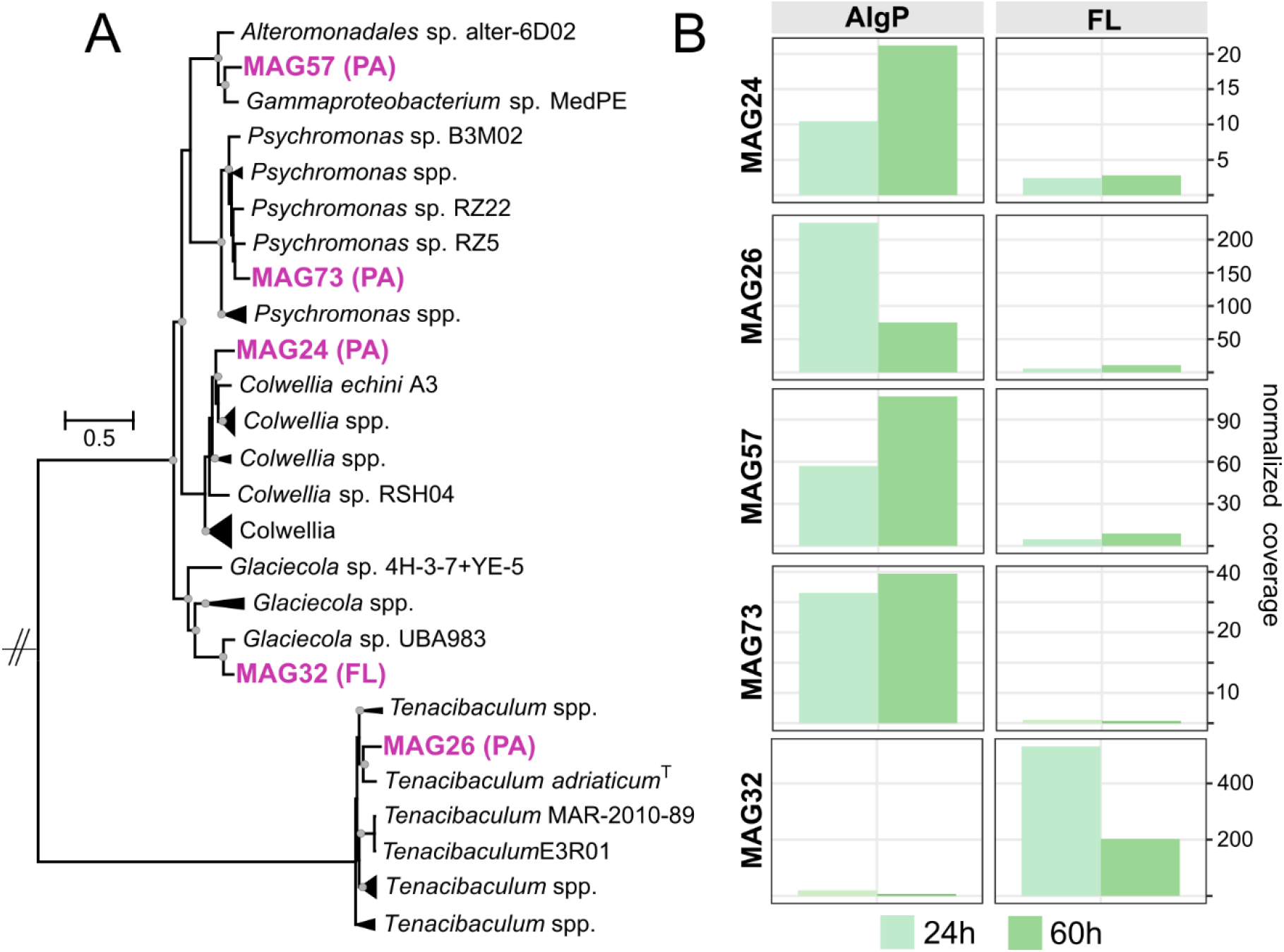
Phylogeny and abundance of metagenome-assembled genomes (MAGs). **A:** Maximum-likelihood phylogeny based on 92 single-copy core genes in context of related genomes. Dots designate nodes with >90% bootstrap support. Supplementary Fig. 5 shows an extended tree including medium-quality MAGs and additional related genomes. **B:** Normalized coverage in metagenomes at 24 and 60 h.

Approximately 3% of genes in the major PA-MAGs were annotated as CAZymes (Table 1), mostly PL6, PL7 and PL18 alginate lyases with CBM32, CBM16 or CBM6 domains (Fig. 5). Considerable transcript abundances of alginate lyase and monomer-processing genes *kdgA*, *kdgF*, *kdgK* and *dehR* (Supplementary Table 4) illustrate the complete metabolization of alginate. Approximately half of CAZymes from *Colwellia*, *Psychrobium* and *Psychromonas* MAGs harbor predicted signal peptides and were hence likely secreted (Supplementary Table 4). Presumably, CAZyme secretion into the polysaccharide matrix facilitated particle utilization, enhancing polymer hydrolysis and subsequent oligomer uptake [91]. MAG32 affiliated with the dominant FL taxon *Glaciecola* encoded only three PLs but eighteen GHs, with highest transcript abundances of families GH3, GH13 and GH23 (Fig. 5). The lower fraction of signal peptides in its CAZymes (30%) indicates that secreted enzymes are less relevant when free-living, pointing towards opportunistic interactions with primary hydrolyzers.

**Fig. 5:**
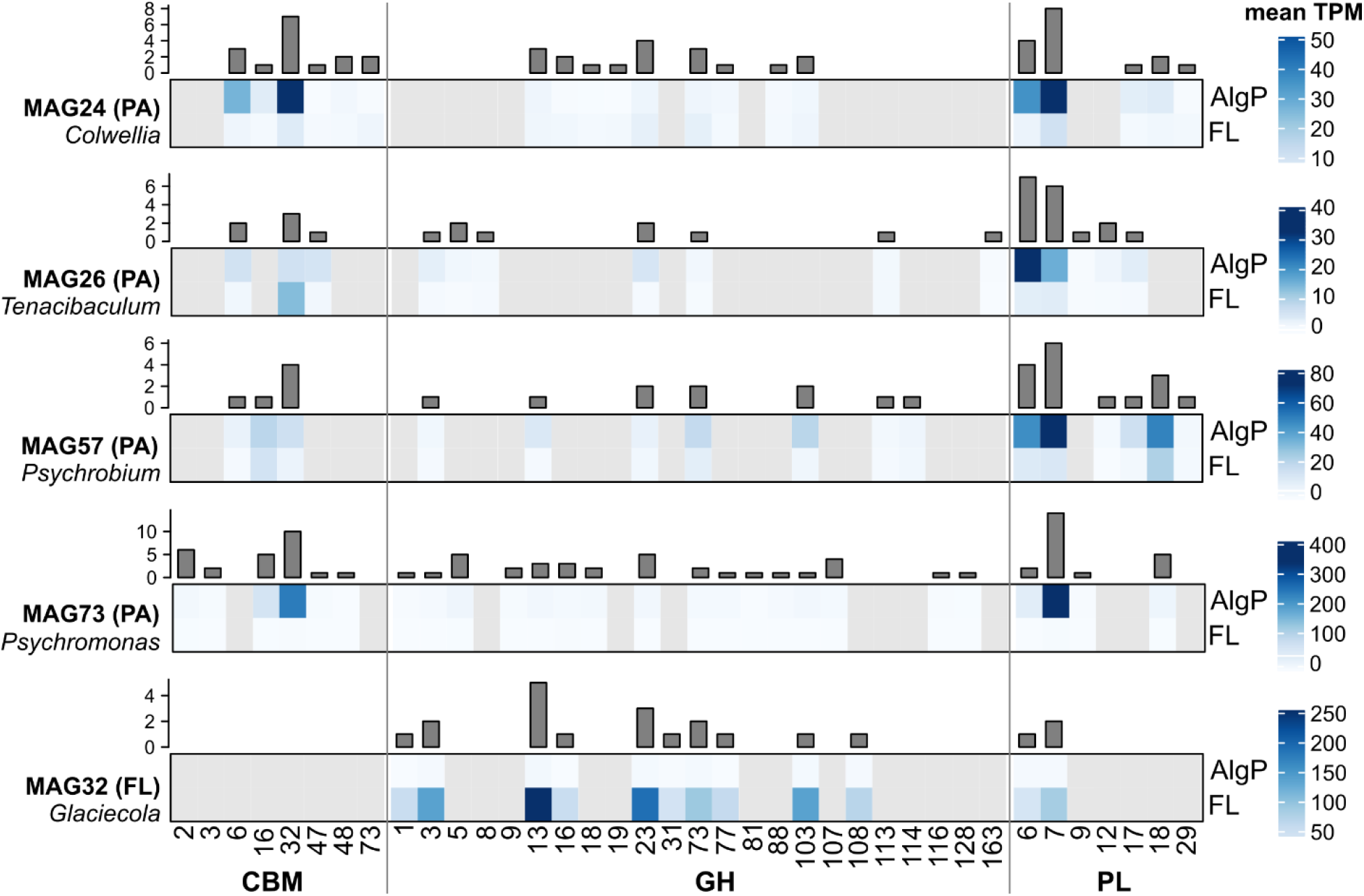
Diversity and expression of CAZymes in metagenome-assembled genomes. Relative transcript abundance (heatmaps) and number (barplots) of CAZyme-encoding genes in MAGs affiliated with the dominant PA and FL community members.

Overall, only some CAZymes of each MAG’s repertoire showed elevated transcript abundances (Fig. 5). We assume that the “silent” CAZymes enable the degradation of other carbohydrates. For instance, *Colwellia*-MAG24 encodes a homolog of the rarely described PL29 family (locus tag 50313), potentially activated in presence of chondroitin sulfate, dermatan sulfate or hyaluronic acid [92]. Although MAG24 clusters with the hydrolytic model isolate *Colwellia echini* A3 (Fig. 4A) at 80% average nucleotide identity, a BLASTp survey revealed that CAZymes targeting agar, carrageenan and furcellaran are not shared with strain A3 (Supplementary Table 4). Divergent CAZyme repertoires in related *Colwellia* spp. presumably reflect their different niches, as *C. echini* has been isolated from a sea urchin [60].

#### MAG-specific polysaccharide utilization loci (PUL)

We detected several PUL in MAG26 affiliated with *Tenacibaculum*. For instance, one PUL encodes PL12 and PL17 alginate lyases plus SusCD transporters, the hallmark of flavobacterial PUL (Fig. 6A). In contrast, CAZyme genes in gammaproteobacterial MAGs were largely scattered throughout the genomes, although PUL-like operons occur in related taxa [60, 93, 94]. *Psychromonas*-MAG73 encodes two PL7 from subfamilies 5 and 3, each harboring a CBM16 and CBM32 domain (Fig. 6B, Supplementary Table 4). Together with the adjacent CBM16 gene, this combination indicates efficient binding and processing of different alginate architectures [95, 96]. *Psychrobium*-MAG57 encodes a PL12, a candidate novel variant of alginate lyases shared with other bacteria from Helgoland [67]. Co-localization of this PL12 with exopolysaccharide-related genes (Fig. 6C) might link polysaccharide degradation and biosynthesis, considering the regulation of exopolysaccharide metabolism via PLs [97, 98]. A comparable gene arrangement in an alginolytic *Maribacter* strain from the south Atlantic [99] supports the potential implications for particle colonization.

**Fig. 6:**
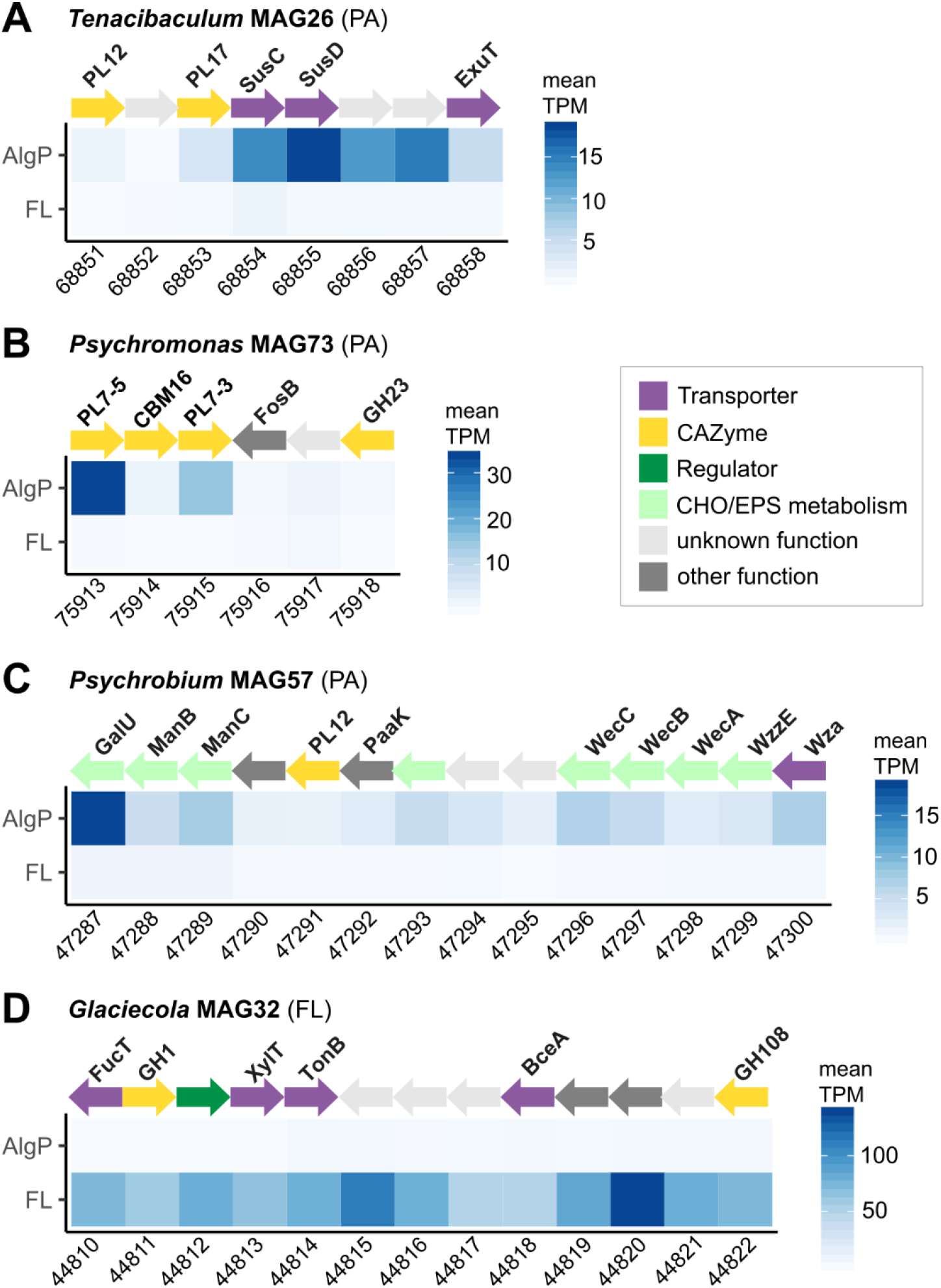
Structure and expression of PUL in MAGs. **A:** PUL encoding SusCD and a PL12-PL17 pair in *Tenacibaculum*-MAG26. **B:** PL7 genes from different subfamilies co-localized with a CBM16 gene in *Psychromonas*-MAG73. **C:** PL12 and exopolysaccharide-related genes (green) in *Psychrobium*-MAG57. **D:** Unique GH108 adjacent to GH1 and carbohydrate transporter genes in *Glaciecola*-MAG32. Supplementary Table 4 and Supplementary Fig. 6 show detailed gene annotations and PUL architectures. EPS: exopolysaccharide; CHO: carbohydrate.

A GH108 gene unique to *Glaciecola*-MAG32, co-localized with GH1 and carbohydrate transporter genes (Fig. 6D), might allow scavenging oligomers released from particles. MAG32 is related to the deep-sea isolate *Glaciecola* sp. 4H-3-7+YE-5 (Fig. 4A), and their sharing of 40 CAZymes including a GH13 pair and adjacent GH77 (locus tags 06167−06169) indicates ecological relevance in diverse habitats [100]. Notably, MAG32 encodes PL6 and PL7 lyases and the entire monomer processing pathway (Supplementary Table 4), indicating the ability for alginate depolymerization. However, considering its lower number of alginate lyase genes than PA-MAGs, MAG32 might be outcompeted on particles. Alternatively, its truncated version of the *A. macleodii* alginate operon (Supplementary Fig. 7A) in combination with *dehR* and *kdgF* being encoded in distant genomic locations (Supplementary Table 4) might signify inefficient or even lost alginolytic capacity.

#### Diversity of PL7 homologs in MAG73

The presence of fourteen PL7 genes in *Psychromonas*-MAG73 signifies a marked specialization towards alginate, as hydrolytic activity scales with CAZyme number [17]. Two of these homologs (locus tags 20477 and 38641) exhibit the highest transcript abundances of all PL7 genes in our dataset (Supplementary Table 4). Both are related to biochemically characterized lyases from *Vibrio* strains (Supplementary Fig. 8) isolated from macroalgae or seawater [101–103]. For instance, PL7_38641 has 72% amino acid identity to AlyD of *Vibrio splendidus*, an endolytic lyase releasing three oligomer fractions especially from guluronate-rich sections [101]. Prediction of a Lipo signal peptide while lacking a CBM indicates that PL7_38641 is anchored as outer membrane lipoprotein (Supplementary Table 4), comparable to AlyA5 from *Zobellia galactanivorans* [104].

The two adjacent PL7 genes from different subfamilies (Fig. 6B) possess predicted Sec and Lipo signal peptides respectively (Supplementary Table 4), indicating complementary membrane-bound vs. secreted localization to maximize alginate utilization. Homologs of PL7_75913 with >50% amino acid identity also occur in *Simiduia* (Cellvibrionales), *Reichenbachiella* and *Marinoscillum* (Cytophagales), indicating wide ecological relevance [105]. PL7_75915 from the poorly described subfamily 3 has 55% amino acid identity to a structurally resolved lyase from *Persicobacter* (Sphingobacteriia) specialized towards alginate of high molecular weight [96], suggesting a role in initial depolymerization. We hypothesize that the two PL variants originate from separate horizontal acquisition events, with subsequent insertion into the same genomic locus. Overall, highly variable transcript abundances of PL7 genes (Supplementary Fig. 8) suggests that different variants are activated by specific biochemical conditions, for instance corresponding to different alginate substructures (e.g. polymer length; dissolved or particulate form; or the ratio of mannuronate and guluronate monomers).

#### Taxa with recurrent occurrence on alginate particles

We compared MAG73 and MAG26 with *Psychromonas* sp. B3M02 and *Tenacibaculum* sp. E3R01 respectively, which were isolated in a comparable study on alginate particles [15]. Supported by ~80% average nucleotide identity and core-gene phylogeny (Fig. 4A), these represent related species with presumably wide ecological relevance on polysaccharide particles. MAG73 and strain B3M02 share nine homologous PLs (Supplementary Table 5), however encoded in different chromosomal contexts. This variable organization, together with the higher PL count in MAG73, indicates CAZyme microdiversity and genomic rearrangements among hydrolytic *Psychromonas*. *Tenacibaculum*-MAG26 and strain E3R01 share 29 homologous CAZymes; however, no polysaccharide lyases were detected in E3R01 while MAG26 encodes seventeen (Supplementary Table 5). These observations indicate CAZyme-related niche specialization among *Tenacibaculum* species, supported by variable PL counts in *Tenacibaculum* type strains (ranging from eight in *T. jejuense* to none in *T. mesophilum* [90]).

#### A single, rare pectin degrader

Despite the compelling evidence that alginate was the preferred bacterial substrate, MAG21 is a candidate for pectin utilization. This medium-quality MAG with 76% estimated completeness accounted for ~95% of differentially abundant transcripts on PecP (Fig. 7A). These patterns corresponded to significantly elevated GH abundances and normalized coverage compared to AlgP (Fig. 7B, Kruskal-Wallis test, *p* < 10^−16^). MAG21 encodes several CAZymes for galacturonate degradation and the entire pathway for processing pectin monomers (Supplementary Table 4). A GH53 gene with the fourth-highest transcript abundance of all CAZymes on PecP (locus tag 118003) presumably cleaves galacturonate-rich side chains from pectinous substrates by endolytic activity [106]. Moreover, the co-localization of GH53 and GH2 plus two carbohydrate transporter genes (Fig. 7B) resembles a galacturonan-related PUL in *Bacteroides thetaiotaomicron* [107]. We hence hypothesize that MAG21 utilizes oligomeric side chains of pectin, processing the resulting galacturonate via tagaturonate and altronate to 2-keto-3-deoxy-D-gluconate through UxuABC (Supplementary Table 4). Alternatively, despite lower transcript abundances, a GH105 gene in another PUL (Fig. 7B) might mediate the exolysis of polymeric pectin, allowing complete saccharification without pectate lyases [87]. The majority of CAZymes in MAG21 possesses homologs in *Catenovulum* spp. (Supplementary Table 4), suggesting that MAG21 is related to this taxon. Accordingly, *Catenovulum* ASVs were only detected on PecP (Fig. 1C).

**Fig. 7:**
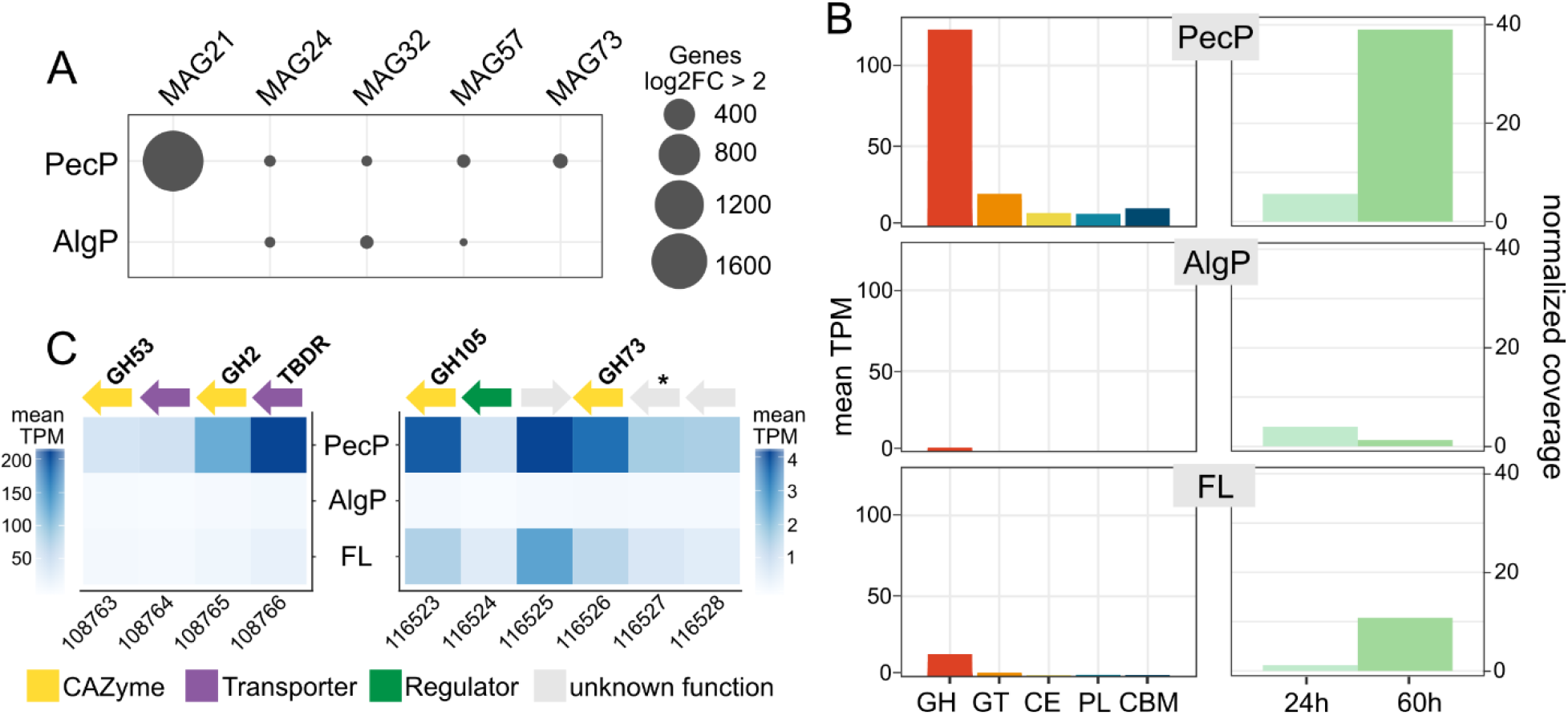
MAG21 as candidate for pectin degradation. **A:** Numbers of differentially abundant transcripts compared to the major PA-MAGs on PecP versus AlgP. **B:** Elevated transcript abundances of glycoside hydrolase genes on PecP. **C, left panel:** PUL with similarities to the pectinolytic operon BT4667−4673 in *Bacteroides thetaiotaomicron*. **C, right panel:** PUL encoding GH105 and GH73 genes plus a hypothetical protein with 60% amino acid identity to an alpha-amylase from *Paraglaciecola arctica* (UniProtKB accession K6ZD77; indicated by asterisk).

An additional PecP-specific pattern occurred in MAG73, with significantly higher transcript abundances of a hybrid biosynthetic gene cluster (Supplementary Table 3, Supplementary Fig. 7B). Homologs of the siderophore-encoding section have been identified in diverse marine bacteria including *Alteromonas* sp. 76-1, with shown iron-chelating activity [108]. Hence, we assume the same functionality in MAG73, presumably to counteract iron limitation on PecP. The upstream spermidine-related section has ~40% amino acid similarity to the polyamine synthesis pathway of *Vibrio*, indicating a PecP-specific role in biofilm formation [109].

## ECOLOGICAL CONCLUSIONS

The predominance of alginolytic pathways demonstrates alginate particles as preferred microbial substrate, while pectin was primarily a colonization scaffold. The establishment of similar communities opposed the hypothesized segregation by substrate preferences. On the contrary, expression of alginate lyases when attached to pectin signify the concept of a “particlescape” including cross-particle interactions. Such a scenario might resemble natural processes when algal polysaccharide exudates enter the water column, self-assemble into particles, and sink to the seafloor. Under such circumstances, bacteria might harvest alginate even if attached to neighboring microhabitats, e.g. by secreting extracellular CAZymes or exploiting hydrolytic activity of co-occurring microbes. The predominance of few taxa indicates that polysaccharide availability stimulates only certain community members, outcompeting most other strains by their extensive CAZyme repertoire. Nonetheless, identification of a single MAG with pectin-specific dynamics indicates that numerically rare but competitive bacteria can establish in specific niches. Altogether, our study illuminates central elements of the biological carbon pump in macroalgae-rich habitats, with implications for microscale ecology, niche specialization and bacteria-algae interactions.

## ACKNOWLEDGMENTS

We are indebted to Antje Wichels, Eva-Maria Brodte, Gunnar Gerdts and Uwe Nettelmann (Biological Department Helgoland) for exceptional scientific and logistic support during the experimental work. Mara Heinrichs, Felix Milke and Mathias Wolterink are thanked for excellent technical assistance. Many thanks to geniaLab (Braunschweig), in particular Ulli Jahnz, for their excellent service and helpful discussions in the preparation of custom polysaccharide particles. Sequencing was carried out by DNASense (Aalborg, Denmark), with professional support by Mads Albertsen, Thomas Yssing Michaelsen, Rasmus H. Kirkegaard and Martin Hjorth Andersen. CB was supported by the HIFMB, a collaboration between the Alfred-Wegener-Institute Helmholtz Centre for Polar and Marine Research and the Carl-von-Ossietzky University Oldenburg, initially funded by the Ministry for Science and Culture of Lower Saxony and VW-Vorab grant ZN3285. HK was supported by Volkswagen Foundation under VW-Vorab grant ‘Marine Biodiversity across Scales’ (MarBAS; ZN3112) and by the Netherlands Organization for Scientific Research (NWO Talent Programme grant VI.Veni.192.086). MW was supported by grant WI3888/1-2 from the German Research Foundation.

## SUPPLEMENTARY FIGURES

**Supplementary Fig. 1.**
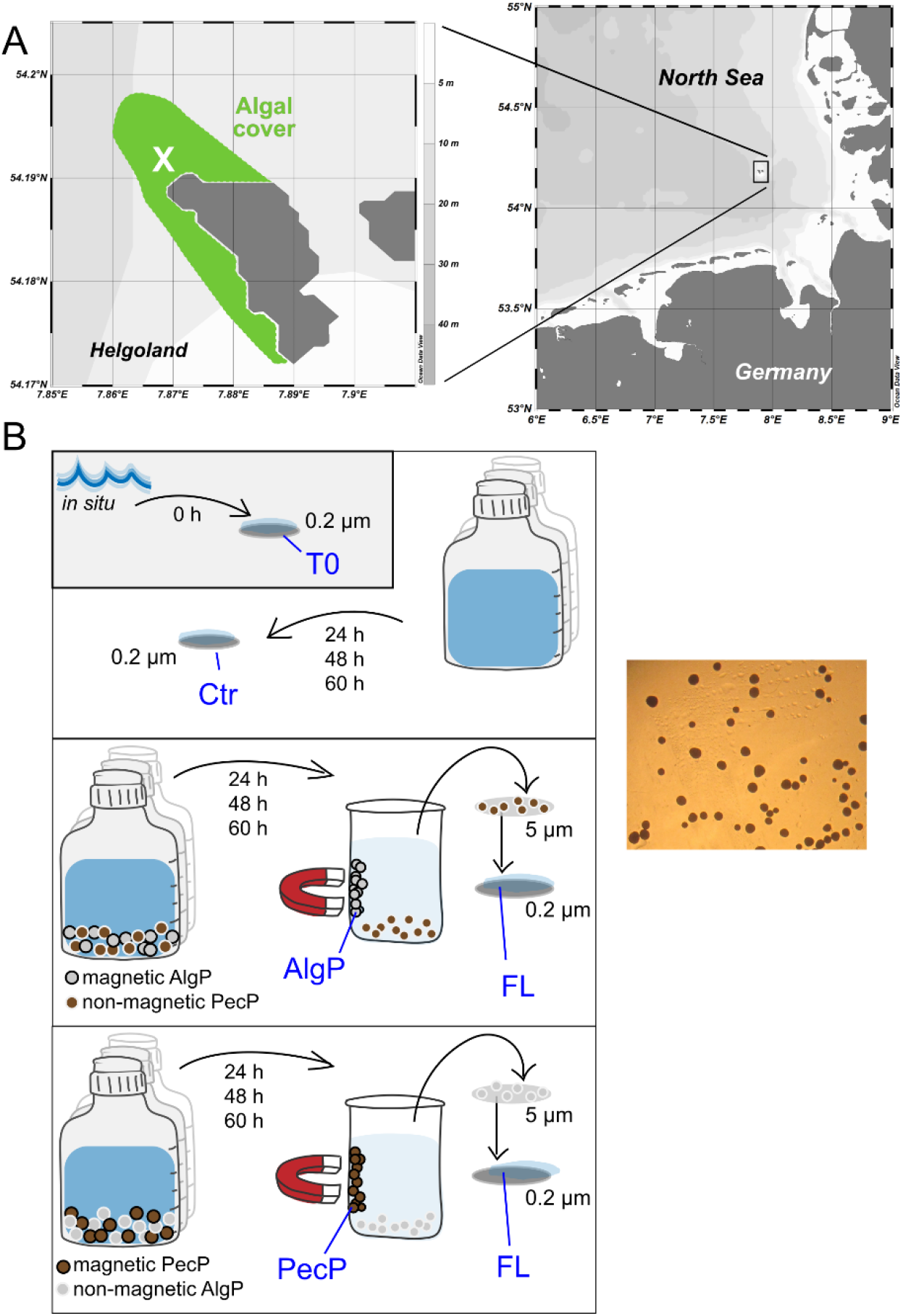
**A:** Sampling location (cross) near Helgoland with dense macroalgal cover; schematically depicted in green. **B:** Experimental setup using polysaccharide particles (insert), allowing separation of magnetic and non-magnetic particles (Supplementary Video). FL communities were sampled by size-fractionated filtration of the supernatant (5 μm followed by 0.2 μm). Ambient seawater (T0) and control samples (CTR) were directly filtered on 0.2 μm. Amplicon (0, 24, 48 and 60h), metagenomic (24 and 60h), and metatranscriptomic sequence data (60h) were generated after different intervals of incubation.

**Supplementary Fig. 2.**
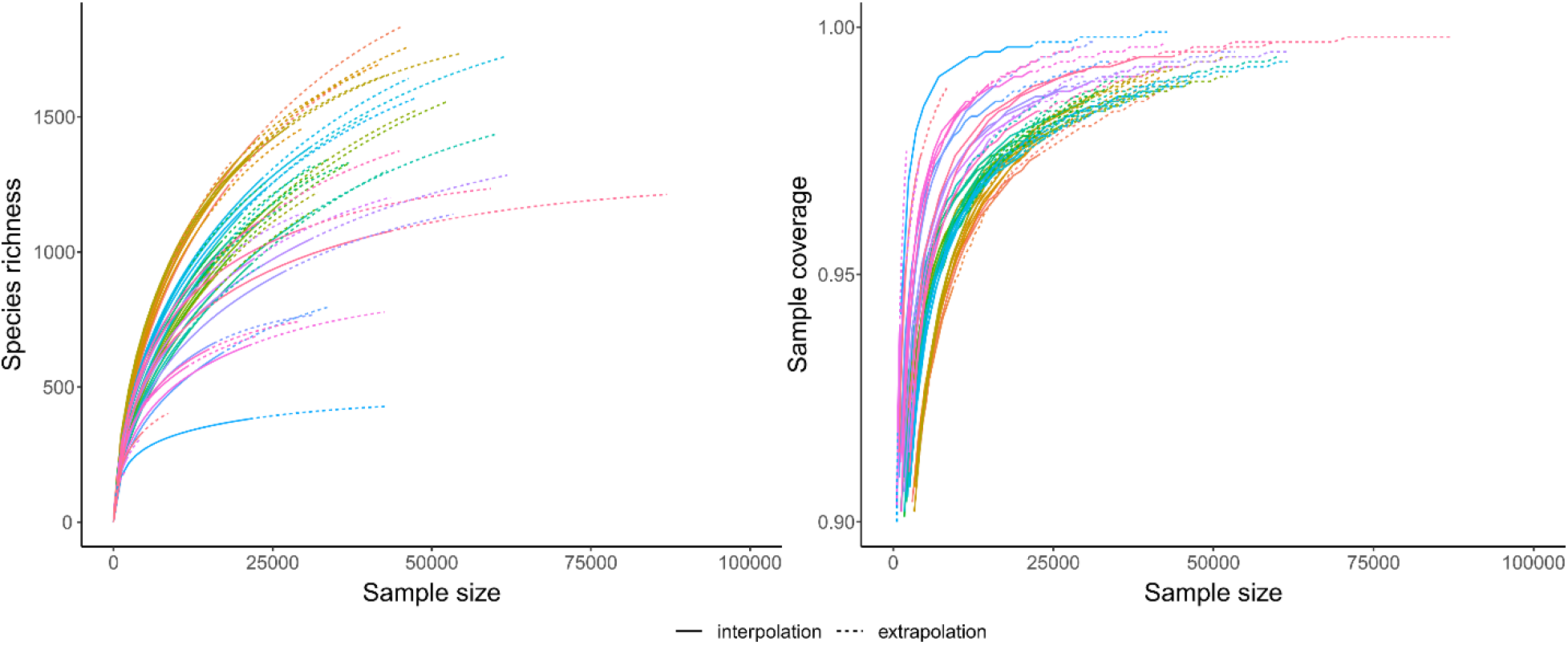
Rarefaction and coverage analysis of amplicon reads.

**Supplementary Fig. 3.**
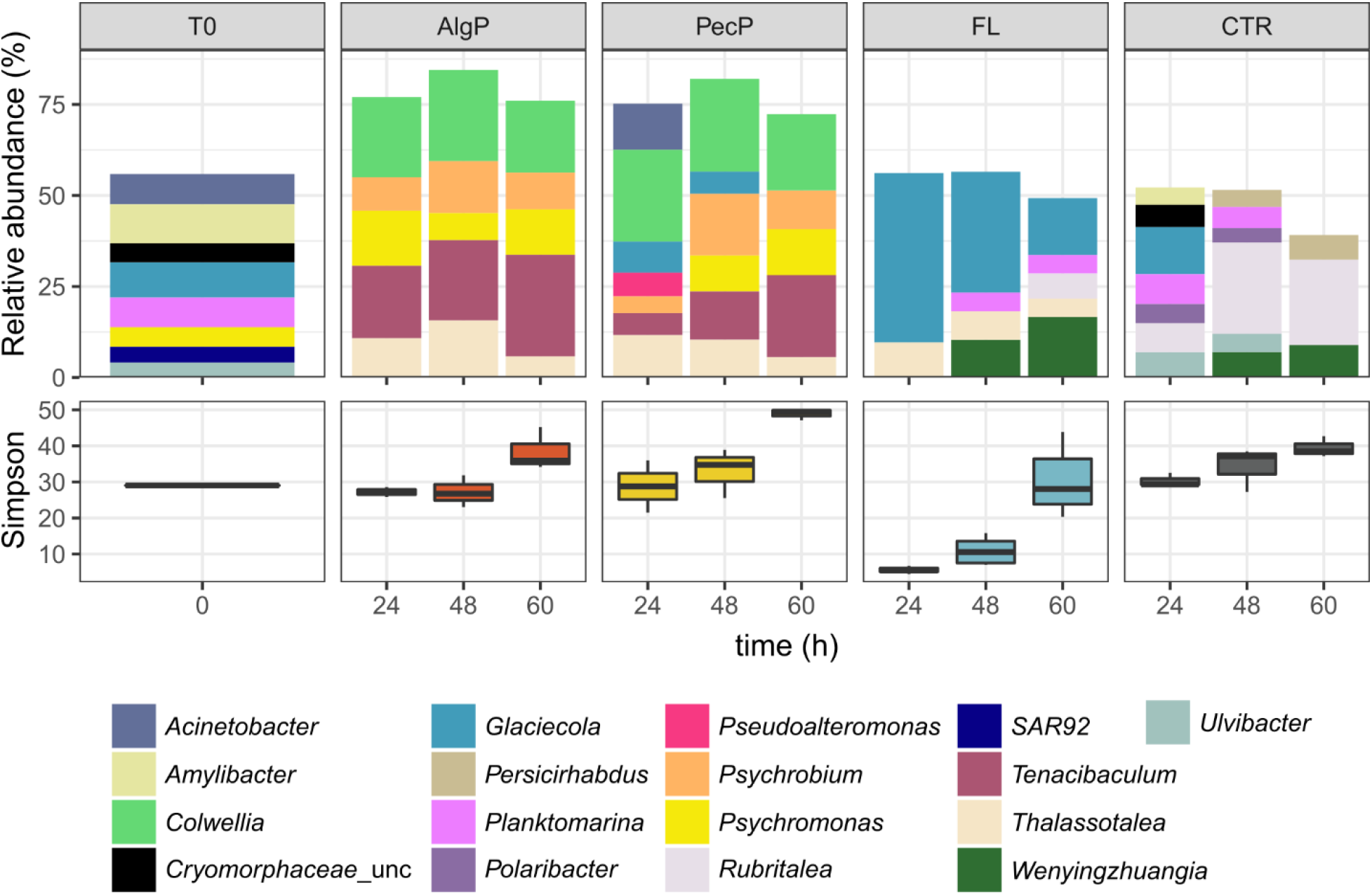
Relative abundances of bacterial genera (top) and bacterial alpha-diversity (bottom) on alginate (AlgP) and pectin particles (PecP), in the free-living fraction (FL), control incubations (CTR) and the ambient community (T0). Only genera with abundances >4% are shown.

**Supplementary Fig. 4.**
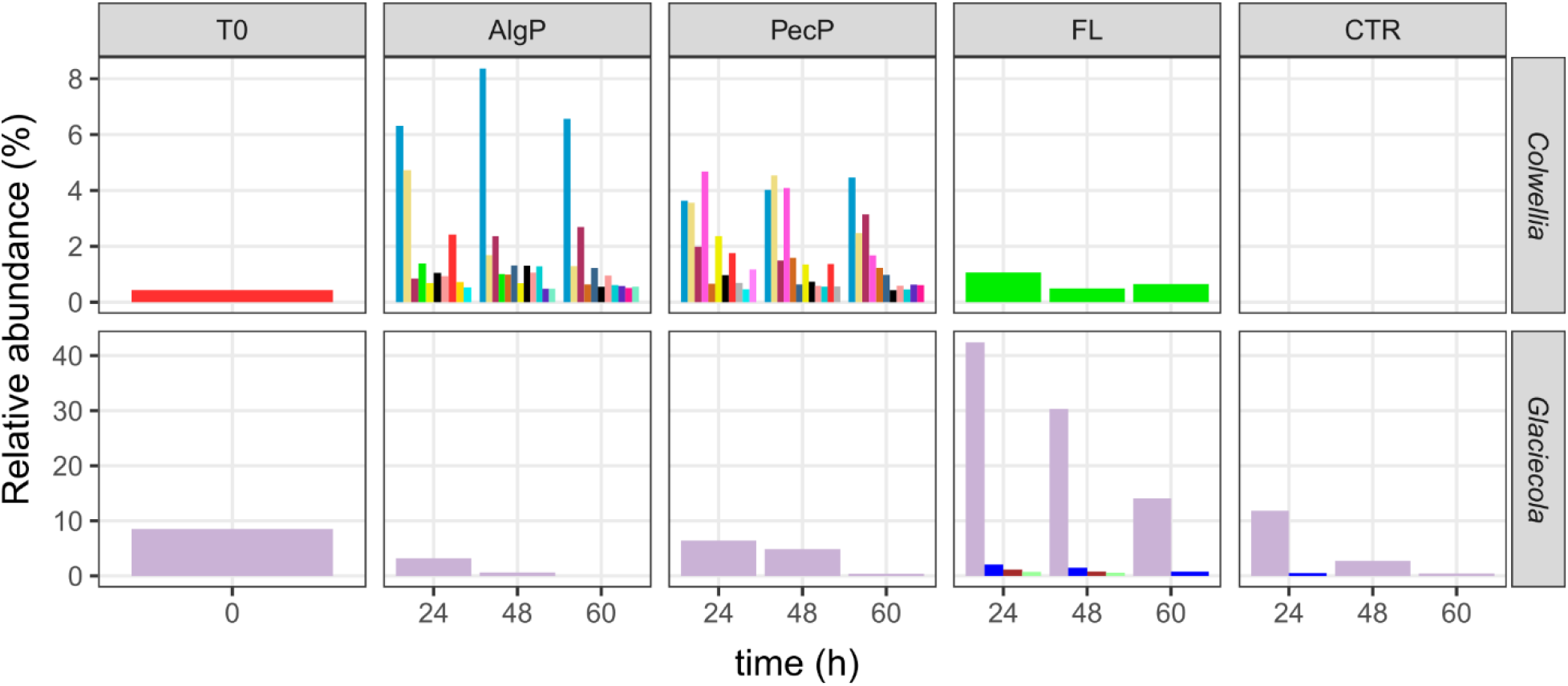
Relative abundances of *Colwellia* and *Glaciecola* ASVs in PA and FL communities. Each color corresponds to a distinct ASV.

**Supplementary Fig. 5.**
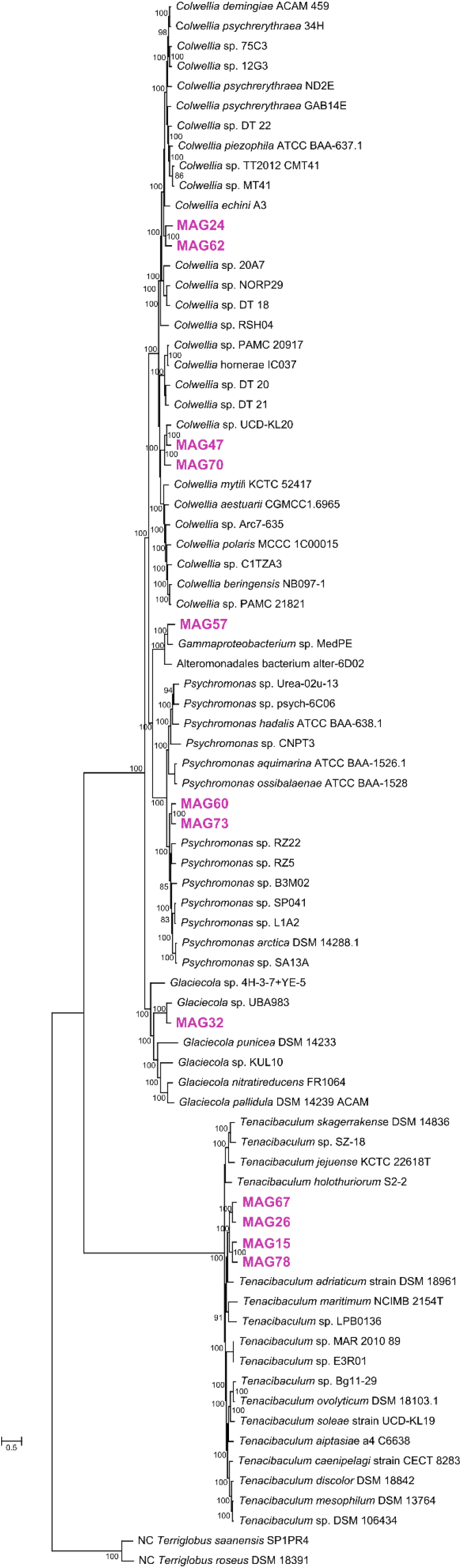
Maximum-likelihood phylogeny of MAGs based on 92 single-copy core genes, including the five near-complete MAGs (>90% completeness, <10% contamination), medium-quality MAGs (>70% completeness, <10% contamination) assigned to the same genus, and other related genomes.

**Supplementary Fig. 6.**
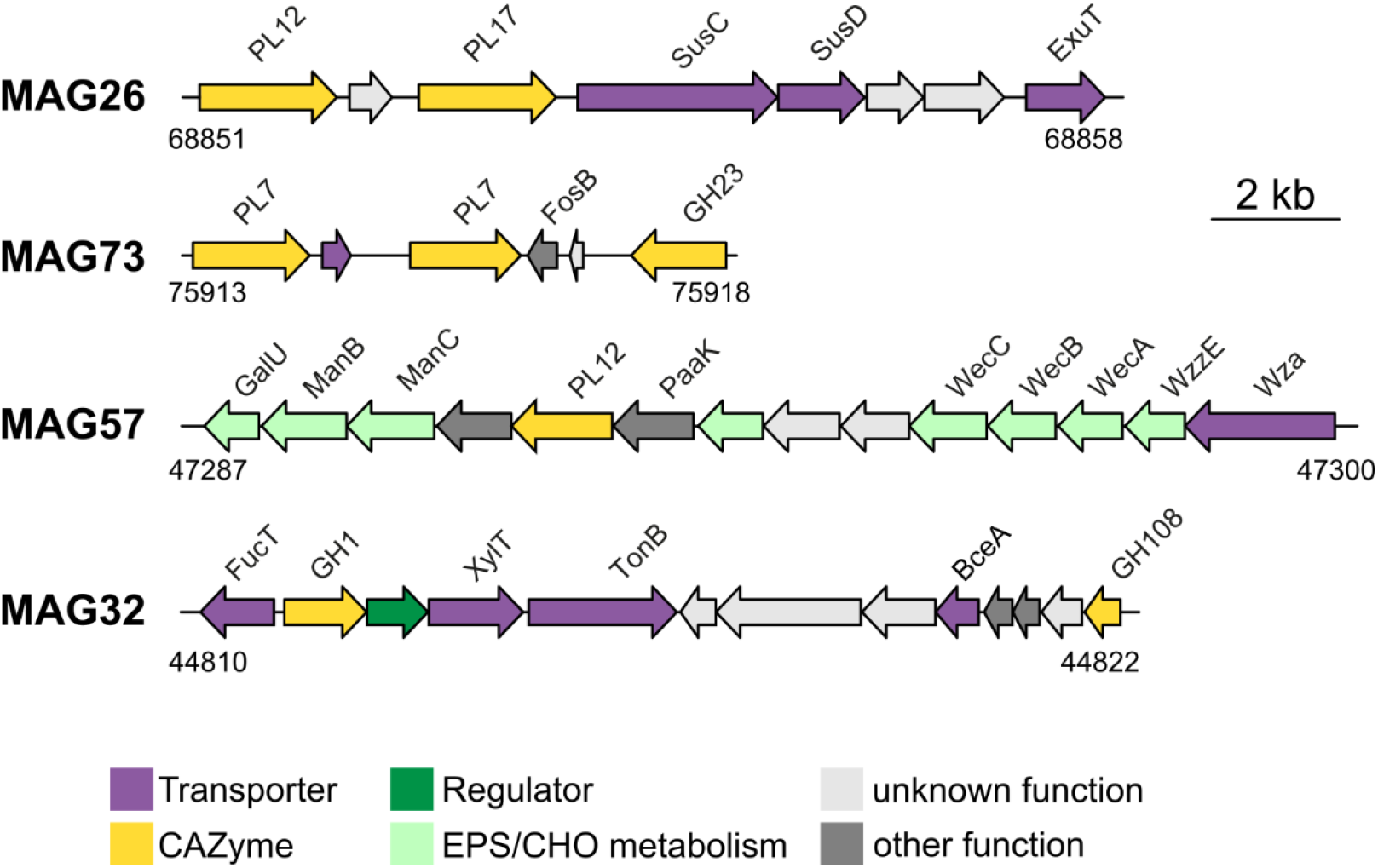
Detailed architecture of PUL in MAGs.

**Supplementary Fig. 7.**
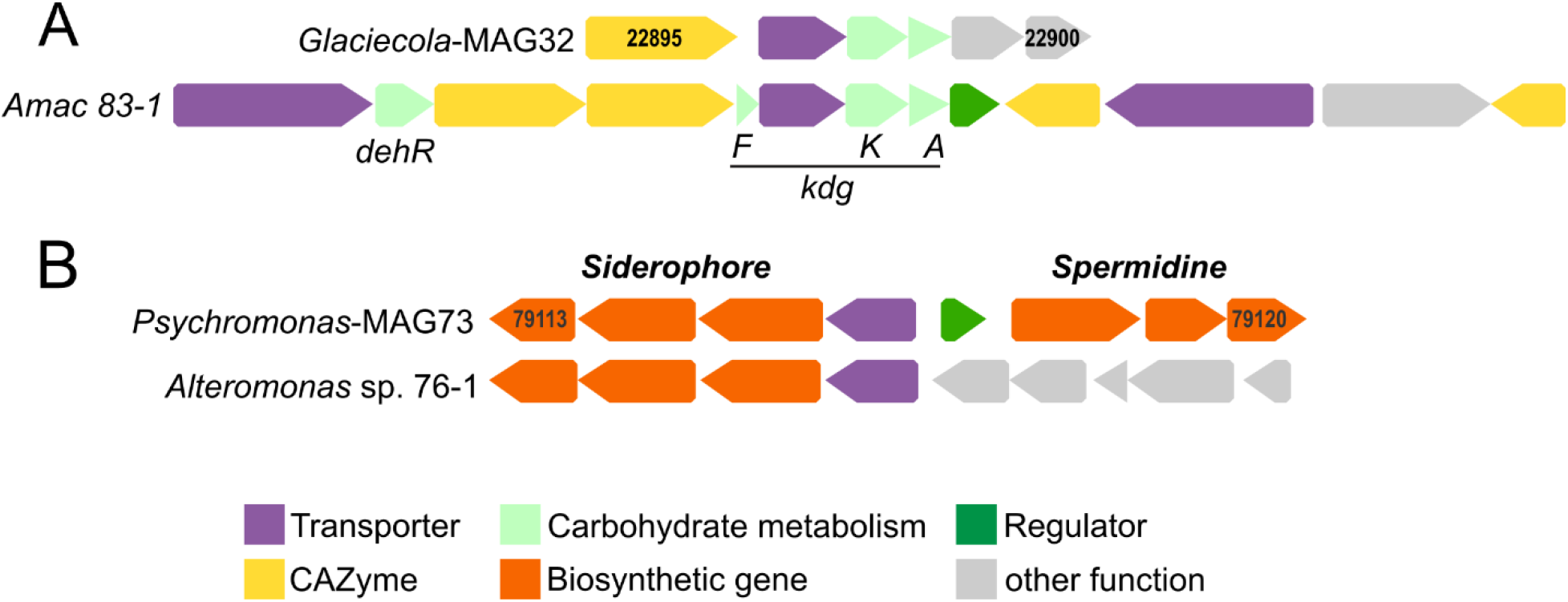
**A:** Truncated alginolytic operon in *Glaciecola*-MAG32 compared to the functionally characterized PUL in *Alteromonas macleodii* 83-1. BLASTp confirmed that missing alginolytic genes were not encoded on another MAG32 contig. **B:** Hybrid biosynthetic gene cluster in MAG73 encoding a siderophore homologous to a functional cluster in *Alteromonas* sp. 76-1 (left section) as well as spermidine-related genes homologous to VC1623 and VC1624 in *Vibrio cholerae* (right section). Locus tags are indicated inside the first and last gene.

**Supplementary Fig. 8.**
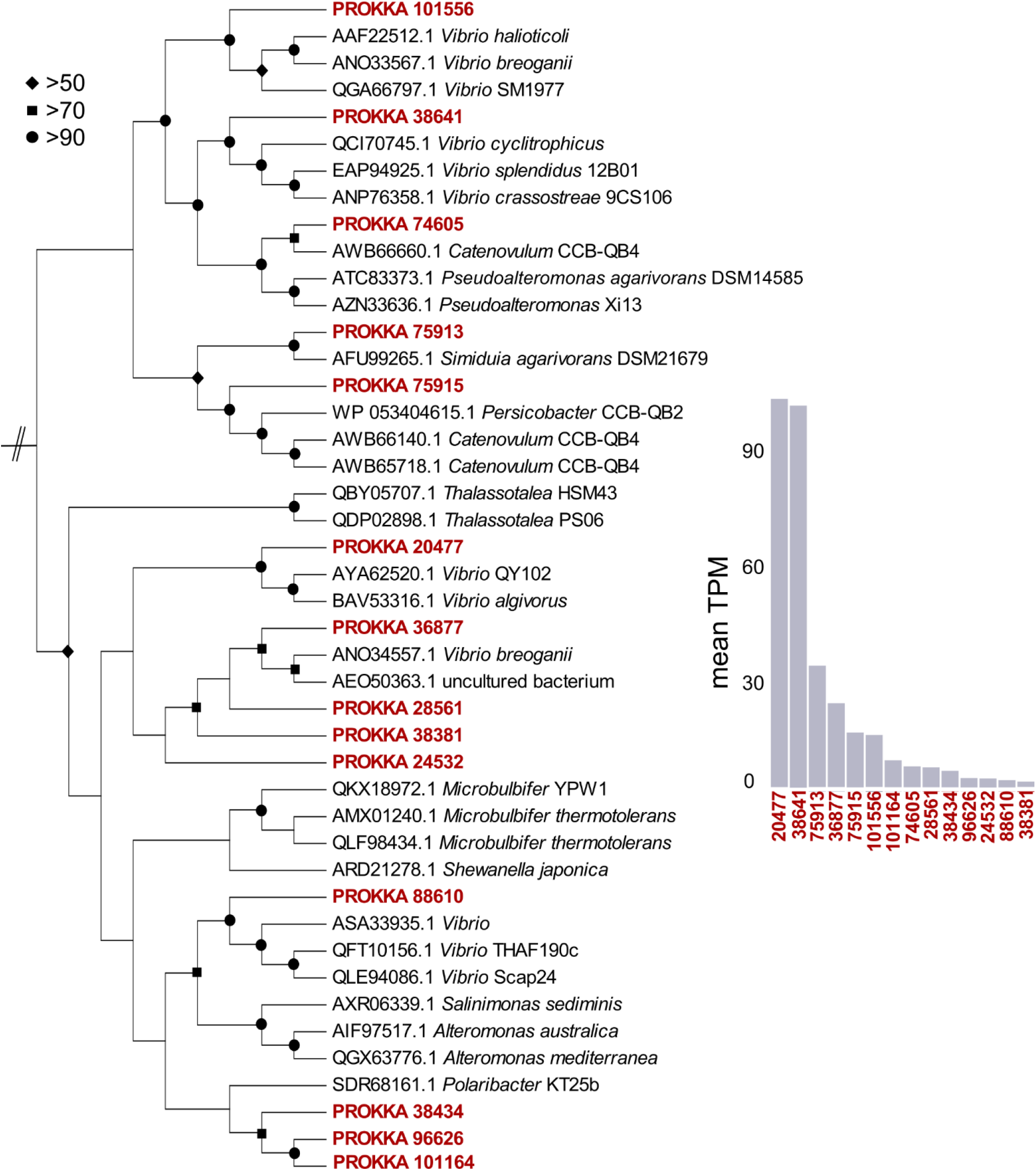
Maximum-likelihood phylogeny of PL7 genes from MAG73, using BAB03312.1 from *Sphingomonas* as outgroup. Only some homologs showed substantial transcript abundances (right insert).

**Supplementary Table 1:** Metadata, sampling strategy, and statistics from amplicon, metagenome and metatranscriptome sequencing.

**Supplementary Table 2:** Complete overview of metagenomic genes, their abundance in metatranscriptomes (transcripts per million), and their assignment to CAZyme and KEGG categories.

**Supplementary Table 3:** Complete overview of differential transcript abundance analysis.

**Supplementary Table 4:** Complete overview of metagenome-assembled genomes.

**Supplementary Table 5:** Comparison of *Tenacibaculum*-MAG26 and *Psychromonas*-MAG73 with *Psychromonas* sp. B3M02 and *Tenacibaculum* sp. E3R01 from Enke et al. 2019 (https://doi.org/10.1016/j.cub.2019.03.047).

**Supplementary Table 6:** Composition of alginate and pectin particles.

**Supplementary Video:** Magnetic polysaccharide particles.

